# Computational Saturation Mutagenesis to predict structural consequences of systematic mutations on protein stability and rifampin interactions in the β subunit of RNA polymerase in *Mycobacterium leprae*

**DOI:** 10.1101/654095

**Authors:** Sundeep Chaitanya Vedithi, Carlos H. M. Rodrigues, Stephanie Portelli, Marcin J. Skwark, Madhusmita Das, David B. Ascher, Tom L Blundell, Sony Malhotra

## Abstract

In contrast to the situation with tuberculosis, rifampin resistance in leprosy may remain undetected due to the lack of rapid and effective diagnostic methods. A quick and reliable method is essential to determine the impacts of emerging detrimental mutations. The functional consequences of missense mutations within the β-subunit of RNA polymerase in *Mycobacterium leprae* (*M. leprae*) contribute to phenotypic rifampin resistance outcomes in leprosy. Here we report *in-silico* saturation mutagenesis of all residues in the β-subunit of RNA polymerase to all other 19 amino acid types and predict their impacts on overall thermodynamic stability, on interactions at subunit interfaces, and on β-subunit-RNA and rifampin affinities using state-of-the-art structure, sequence and normal mode analysis-based methods. A total of 21,394 mutations were analysed, and it was noted that mutations in the conserved residues that line the active-site cleft show largely destabilizing effects, resulting in increased relative solvent accessibility and concomitant decrease in depth of the mutant residues. The mutations at residues S437, G459, H451, P489, K884 and H1035 are identified as extremely detrimental as they induce highly destabilizing effects on the overall stability, nucleic acid and rifampin affinities. Destabilizing effects were predicted for all the experimentally identified rifampin-resistant mutations in *M. leprae* indicating that this model can be used as a surveillance tool to monitor emerging detrimental mutations conferring rifampin resistance in leprosy.

**AUTHOR SUMMARY:** Emergence of primary and secondary drug resistance to rifampin in leprosy is a growing concern and poses threat to the leprosy control and elimination measures globally. In the absence of an effective *in-vitro* system to detect and monitor phenotypic rifampin resistance in leprosy, most of the diagnosis relies on detecting mutations in the drug resistance determining regions of the *rpoB* gene that encodes the β subunit of RNA polymerase in *M. leprae*. Few labs in the world perform mouse food pad propagation of *M. leprae* in the presence of drugs (rifampin) to determine growth patterns and confirm resistance, however the duration of these methods lasts from 8 to 12 months making them impractical for diagnosis. Understanding molecular mechanisms of drug resistance is vital to associating mutations to clinical resistance outcomes in leprosy. Here we propose an *in-silico* saturation mutagenesis approach to comprehensively elucidate the structural implications of any mutations that exist or can arise in the β subunit of RNA polymerase in *M. leprae*. Most of the predicted mutations may not occur in *M. leprae* due to fitness costs but the information thus generated by this approach help decipher the impacts of mutations across the structure and conversely enable identification of stable regions in the protein that are least impacted by mutations (mutation coolspots) which can be a choice for small molecule binding and structure guided drug discovery.

## INTRODUCTION

Nonsynonymous mutations in genes that encode drug targets in mycobacteria can induce structural and consequent functional changes leading to antimicrobial resistance, the burden of which is rapidly increasing and is a global health concern. Diagnosis of ~600,000 new cases of rifampin-resistant tuberculosis in 2018 suggest that it poses a risk for the concomitant increase in undiagnosed rifampin-resistant leprosy worldwide [1]. *Mycobacterium leprae* (*M. leprae*), the causative bacilli for leprosy, is phylogenetically closest to *Mycobacterium tuberculosis* [2] and developed resistance to rifampicin before the introduction of WHO multi-drug therapy (MDT). Despite the long duration of chemotherapy with MDT (six months in paucibacillary to 12 months in multibacillary disease), rifampin-resistant case numbers are less and represent only 3-5% of total relapsed leprosy cases reported in 2017 [3]. One of the possible reasons for the low numbers of drug-resistant leprosy cases worldwide is the lack of quick, effective and reliable *in vitro* diagnostic tests for confirming phenotypic resistance. Current methods rely on identifying drug resistance mutations in *rpoB* gene through gene sequencing and/or to test growth patterns of *M. leprae* in response to drugs in an *in vivo* system (footpads of mice), the later technique is both time and labour intensive.

While mutations within the β-subunit of RNA polymerase contribute to clinical resistance to rifampin, the associated structural changes can complicate the transcription process in bacteria by modulating various complex physiological processes[4], the knowledge of which is essential for novel drug discovery or alternative therapies to treat rifampin resistant strains of *M. leprae*. In the absence of an artificial culture system to propagate and study the molecular mechanisms of resistance, it is exceptionally challenging to define an experimental phenotype for rifampin resistance in leprosy. *M. smegmatis* as a surrogate host with electroporated *M. leprae rpoB* gene has proved a dependable model to study phenotypic effects; however, this technique is limited to biosafety level-2 laboratories that have facilities for gene cloning and sequencing, and cannot be translated to a regular diagnostic setting in leprosy endemic countries[5]. A plausible association between mutations in drug targets and phenotypic resistance outcomes could be established if minimum inhibitory concentrations (MICs) of the drugs are known for the mutant strains. While MICs can be estimated in cultivable species like *M. tuberculosis* and *M. smegmatis*, obtaining growth information from *in vivo* propagation for a slow growing and obligate pathogen like *M. leprae* is often challenging and needs time and resources. *In siiico* methods to predict structural implications of mutations will be extremely useful in understanding mechanisms of drug resistance and help prioritise mutations that require experimental validation in leprosy in the absence of a tool for quantitative estimation of the phenotypic resistance outcomes [6]. Mutations contribute to disruption of protein-ligand and protein-nucleic acid interactions resulting in drug resistance in mycobacterial diseases (Portelli *et al*., 2018; Karmakar *et al.*, 2018). Changes in affinity for the ligand can result from both orthosteric and allosteric mechanism leading to various resistance phenotypes(Vedithi *et al.*, 2018).The β-subunit of RNA Polymerase in *M. leprae* is encoded by the *rpoB* gene (ML1891) whose product is 1178 amino acids in length. The rifampin resistance determining region (RRDR) is located between residue positions 410 and 480. Approximately 40 mutations have been reported in the *rpoB* gene of *M. leprae* that cause clinical resistance to rifampin in leprosy[9–11]; however, in tuberculosis, nearly 270 mutations have been reported in the same gene that shares 96% identity with that of *M. leprae* [12]. As the burden of rifampin resistance is very high in *M. tuberculosis* with known and new mutations being reported from different studies[13–17], it is important to monitor the emergence of new rifampin-resistant mutations in *M. leprae*. A comprehensive understanding of the effects of any mutation on the structure of RNA Polymerase (RNAP) is important in the context of monitoring emerging rifampin resistance and its implications on controlling global leprosy incidence.

In order to decipher the effect of systematic mutations on the stability of the protein structure, protein sub-unit interfaces, nucleic acid and ligand interacting sites, we performed *in-siiico* saturation mutagenesis and predicted the stability changes in protein-protein, protein-ligand and protein-nucleic acid affinities. Additionally, we also assessed the impacts of mutations on the secondary structures of the polypeptide chains, on the relative sidechain solvent accessibility, depth and on the residue-occluded packing density. Residue evolutionary conservation scores were determined and compared with the predicted destabilizing effects. Extremely detrimental mutations were selected and analysed for changes in their interatomic interactions that might explain the destabilizing effects. To explore further the vibrational entropy and enthalpic changes of flexible conformations we employed an empirical force field-based method - FoldX[18], a course-grained normal mode analysis (NMA) based elastic network contact model - ENCoM [19] and a consensus predictor that integrates normal mode approaches with graph-based distance matrix in the mutating residue environment–DynaMut [20]. Finally, fragment hotspots [21] were mapped on the structures to provide information on potential druggable sites whose stability is predicted to be least likely affected by mutations (no mutations in these regions were identified in leprosy). We termed these sites as “Mutation coolspots” which can be explored for novel/alternative small molecule binding and structure-guided drug discovery to treat rifampin-resistant leprosy.

## MATERIALS & METHODS

### Design

The key stages in the methodology involve developing of a model based on the known structures of homologues, quality assessment, generating mutation lists and sequential submissions to stability change prediction servers for sequence, structure and vibrational entropic terms (Fig 1A).

**Fig 1:**
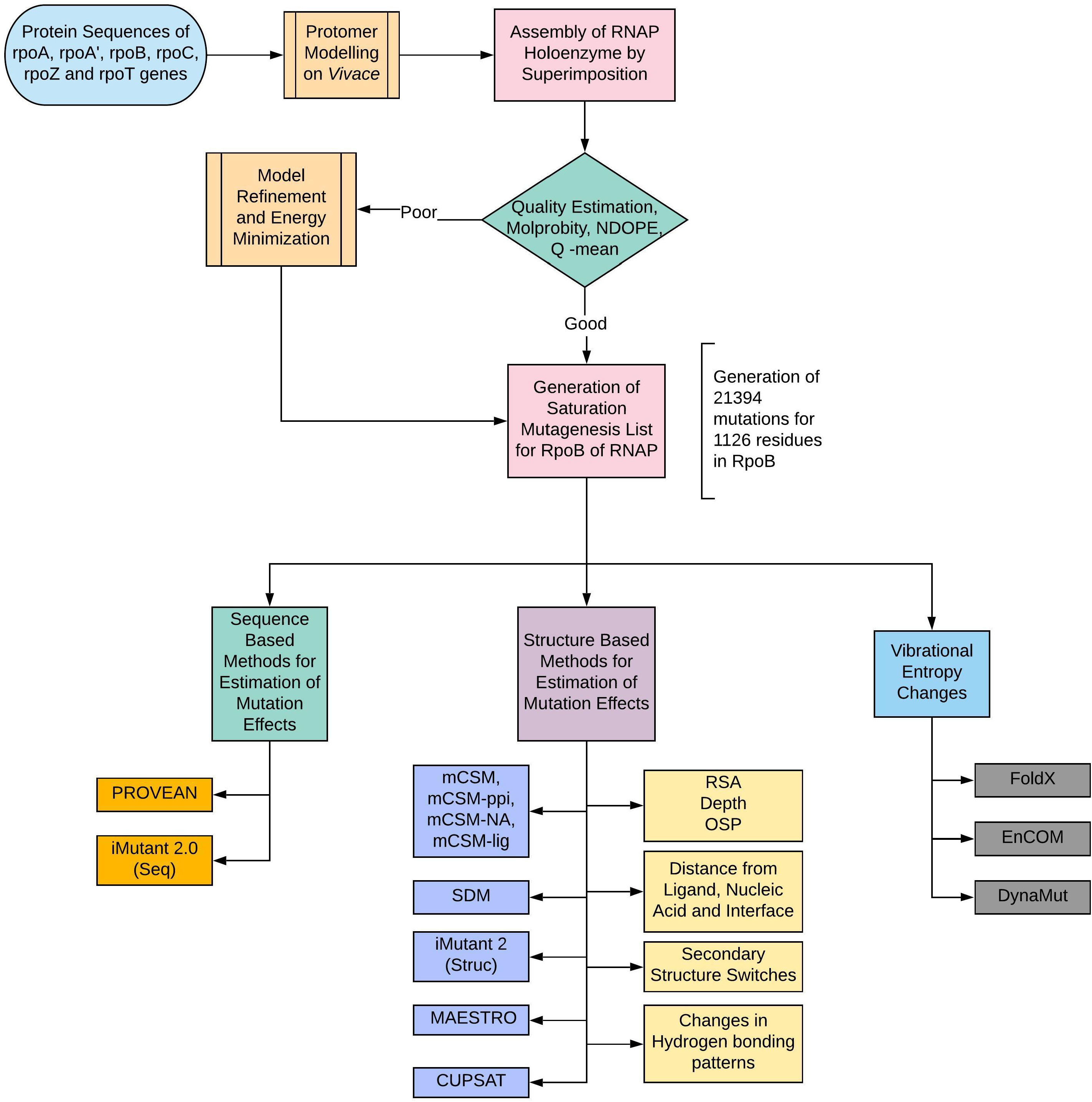
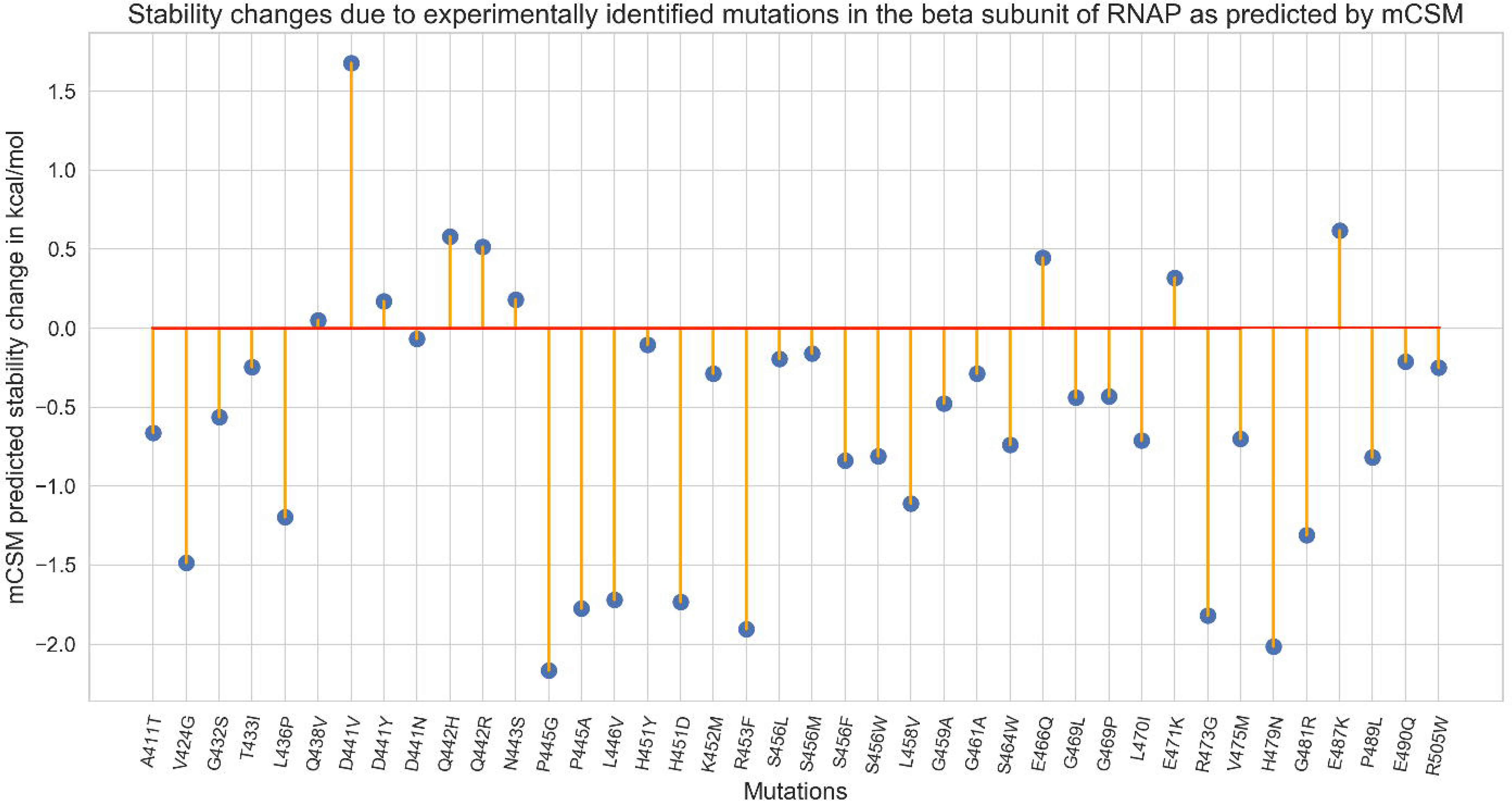
**[A]** Methodology and study design. [B] A lollipop plot with stability predictions for mutations reported in literature known to confer rifampin resistance in Leprosy.

### Comparative modelling, quality assessment and model refinement

A model for RNAP holoenzyme of *M. leprae* was built using Modeller 9.21 based using templates from *M. tuberculosis* (PDB ID:5UH5 (96% identity, 3.8Å resolution) containing RNAP and nucleic acid scaffold with DNA and three nucleotides of RNA complementary to the template DNA strand and PDB ID: 5UHC (96% identity, 4.0Å resolution) containing all the elements similar to 5UH5 and rifampin) as described earlier by us [4]. The quality of the generated model was assessed using Molprobity [22] and atomic clashes were removed by minimizing the energy of the model by 100 steps using Steepest Decent (step size = 0.02 Å) and by 10 steps (step size = 0.02A) using conjugate gradient algorithms. Energy minimizations were performed using UCSF Chimera[23], The mutant models were generated using Modeller 9.21 [24] (mutate_model.py) and sidechains of the mutants were optimized using ANDANTE [25], a program that uses χ angle conservation criteria to optimize the sidechain rotamers.

### Saturated Mutagenesis

A systematic list of 21,394 mutations for residues from P28 to E1153 in the β-subunit (the modelled region) was generated. This list was programmatically submitted to a set of servers that predict protein stability and stability of protein-protein, protein-nucleic acid and protein ligand affinity upon mutations. We also used physics-based potentials to determine impacts of mutations on the RNAP complex in flexible conformations.

### Residue Conservation

Conservation scores for each of the residues in the wild-type model were estimated using CONSURF [26] – a server that uses evolutionary patterns of amino acids/nucleic acids from the multiple sequence alignment and develops a probabilistic framework to calculate evolutionary rates for each residue in the sequence.

### Effects of mutations on Protein Stability and Interactions

The effect of mutations on thermodynamic stability of the protein was analyzed using mCSM [27], SDM [28] and FoldX [29]. For SDM, mutant-protein models were generated using ANDANTE [25], an in-house-developed software that considers conserved χ angle conservation rules while identifying the most probable sidechain rotamers for the mutant residues. The effect of mutations on RNA affinity is assessed using mCSM-NA2[30] on mutant models with nucleic acid scaffold. The holoenzyme complex of RNAP consists of five subunits and the effects of mutations on the protein-protein interfaces (between β and all the other sub-units in RNAP complex) were assessed using mCSM-ppi. Rifampin binds to the β-subunit of RNAP and we analyzed the effects of mutations on the protein-ligand affinity using mCSM-lig [31]. Only residues within 10Å of the interatomic distance to rifampin were analyzed by mCSM-lig.

The stability changes were further compared with predictions from other sequence-[PROVEAN[32], I-Mutant 2.0 (Sequence)[33] and structure-based (MAESTRO[34], CUPSAT[35], I-Mutant 2.0 (Structure)) computational tools in order to estimate the reliability of the predictions.

### Changes in Vibrational Entropy and Normal Mode Analysis

In order to determine the effects of the mutations in flexible conformations on protein stability, we used FoldX [18], an empirical force field approach that calculates free energy changes between native and mutant forms of the protein, and an elastic network contact model (ENCoM)[19], which is a coarse grain NMA method that considers the nature of the amino-acids and aids in calculating vibrational entropy changes upon mutations. We also used DynaMut [36], a consensus predictor of protein stability based on the vibrational entropy changes predicted by ENCoM and the stability changes predicted by graph-based signature approach of mCSM.

### Conformational Changes

Conformational changes and their impacts on biophysical properties of the proteins were estimated using SDM [28]. The interatomic distances between each residue and the interface with other subunits in the RNAP holoenzyme, rifampin and nucleic acids in the structure were measured and included in the analysis. Secondary structure switches in mutants, changes in relative solvent accessibility, depth of the residue in Å and residue-occluded packing densities were determined for all the mutations.

### Interatomic Interactions

A few mutations that were experimentally validated elsewhere and are known to be extremely detrimental to stability and ligand interactions were selected and changes in interatomic interactions of the mutating residues were documented using Arpeggio[37], a program that maps the types of interatomic interactions wildtype and mutant residues with the environment based on atom type, interatomic distance and angle constraints. A set of mutations that are not experimentally identified but computationally predicted to have detrimental effects were also chosen from the saturation dataset and a similar analysis was performed. Intermezzo (Bernardo Ochoa Montano & Blundell TL unpublished) was also used for interactive analysis of bonding patterns on Pymol sessions.

### Fragment Hotspot Maps

Fragment hotspot maps [21] aid in locating specific sites on the surface of the protein that are topologically, chemically and entropically favorable for small molecule (fragment) binding. The atomic hotspots on the drug target are explored computationally using donor, acceptor and hydrophobic fragment probes, and introducing a depth criterion to assist in estimating the entropic gain in displacing “unhappy” waters. For ligand-binding proteins, the fragment hotspot maps aid in understanding the pharmacophore characteristics of the interacting regions. We mapped the hotspots on the β-subunit of RNAP and colored the surface with regions that are least impacted by any mutations (mutation coolspots).

## RESULTS

In total 21,394 mutations were generated from 1126 amino acid residues in the β-subunit of RNAP (Supplementary Table-1). The list of experimentally identified mutations and their effects are separately shown in Supplementary Table-2.

### Multivariate analysis of free energy changes predicted by different computational tools for saturated mutations

Along with the in-house developed mCSM and SDM tools for prediction of protein stability changes upon saturated mutagenesis of the β-subunit of RNAP, a comparative analysis was performed with other sequence (PROVEAN, I-mutant 2.0 - Sequence), structure-(CUPSAT, I-mutant 2.0-structure, MAESTRO) and NMA-based tools (FOLDX, ENCOM, DynaMut). Average stability changes caused by all possible mutations at each residue position in the β-subunit of RNAP, as predicted by mCSM and SDM, were compared with other structure-based predictors (Supplementary Fig 1) (rifampin-interacting residues are highlighted). Correlation of overall stability predictions performed by mCSM with each of the other tools indicated an “r” value of 0.55 with SDM, 0.61 with MAESTRO, 0.72 with Imutant 2.0 (Structure) and 0.43 with CUPSAT. Correlations between mCSM, SDM and other sequence and NMA based tools are shown in supplementary figures 2 and 3. The rationale for performing these correlations is to understand how mCSM and SDM being structure-based predictors of stability changes upon mutations, relate to sequence-based methods and vibrational entropy changes in normal mode perturbations.

### Experimentally Identified Mutations

We performed a systematic literature review to list all the mutations reported in the β-subunit of RNAP in *Mycobacterium leprae*. We noted 40 mutations at 32 unique residue positions. The reference articles are listed in Supplementary Table-2. As depicted in Fig 1B, 77.5% (31) of the experimentally identified mutations destabilize the β-subunit. Except for A411T and V424G mutations, all the other residues are present in close proximity to rifampin binding sites (Fig 2A) and destabilize rifampin interactions (mCSM-lig).

**Fig 2:**
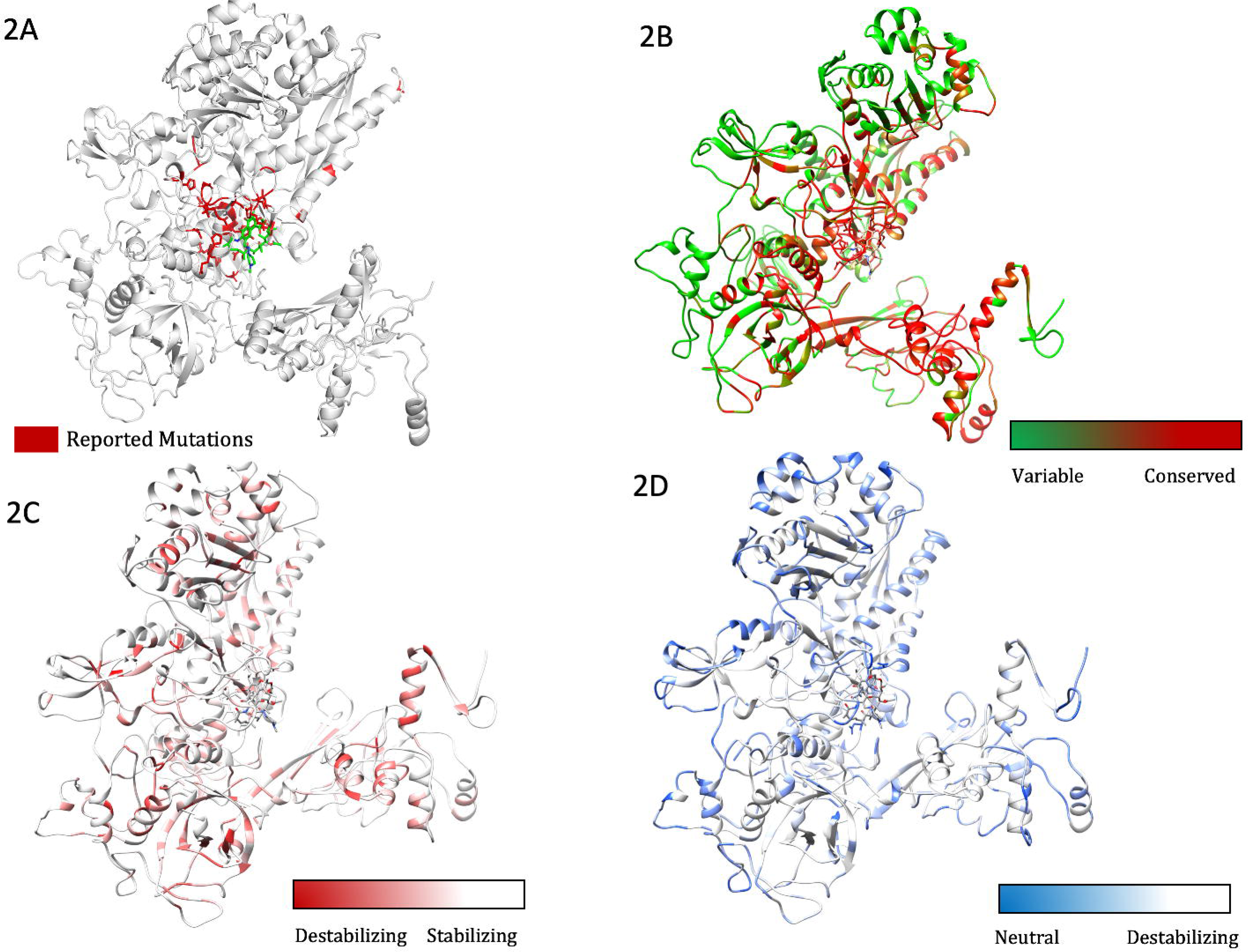
**[A]** The β subunit of RNAP with residues where mutations were reported experimentally from patient samples in various studies (highlighted in red). **[B]** Each residue in the β subunit of RNAP that is colored based on the conservations scores of CONSURF. The residues in green are variable (conservations scores greater than 1) and are usually surface exposed. The residues in red are conserved with conservation scores less than 1 and usually form the core of the protein. The rifampin binding site is highly conserved in *M. leprae*. **[C]**The maximum destabilizing effects (predicted by mCSM) on the protein stability, a mutation can induce at each residue position, is mapped on the structure. Red are the regions that are largely destabilized by mutations while the white regions are relatively stable with mutations. **[D]** The converse of B where the regions that are predicted to be least impact the stability with any mutation are coloured in blue and we called them “Mutation CoolSpots”.

### Residue conservation and protein stability

The stability changes, predicted after saturation mutagenesis of each residue in the β-subunit, were compared with residue conservation scores. CONSURF scores of less than zero are attributed to conserved residues [26] and scores of zero and above to variable residues (score 3 being maximum and highly variable). The average change in protein stability that was predicted by mCSM for mutations at each residue position ranged from 0.823 to −3.033 kcal/mol and that of SDM varied from 2.167 to −4.36kcal/mol. Residues that line the active center cleft and interact with rifampin and the nucleic acid scaffold are highly conserved, while surface exposed residues have variable conservation scores (Fig 2B). Rifampin-interacting residues between residue positions ~400-500 are highly conserved and 87.3% of the saturated mutations in this region destabilize the protein (Supplementary Table 1). The maximum destabilizing effect of mutations at each of these residues varied between −0.311 to −4.311kcal/mol(mCSM). The average destabilizing effect predicted by mCSM for all possible mutations at each residue was mapped on to the structure to identify regions are largely impacted by mutations (Fig 2C). Conversely, the residues whose stability is least impacted by all possible mutations are colored in blue to identify “mutation coolspots” that are potentially areas of choice for targeting with small molecules in drug discovery (Fig 2D).

As part of the RNAP holoenzyme complex, the β-subunit interacts with other subunits and has large interfacial regions. The impact of mutations on the stability of these interfaces was measured using mCSM-ppi. It was noted that the maximum destabilizing effect by any mutation at a particular residue in the interface between β and β’ subunits has an affinity change that ranged from −0.021 to −5.108 kcal/mol (−5.108kcal/mol was noted for mutation W1074R which is not reported experimentally in rifampin resistant leprosy cases). The interfacial region and the stability changes are mapped on the structure (Fig 3A and B).

**Fig 3:**
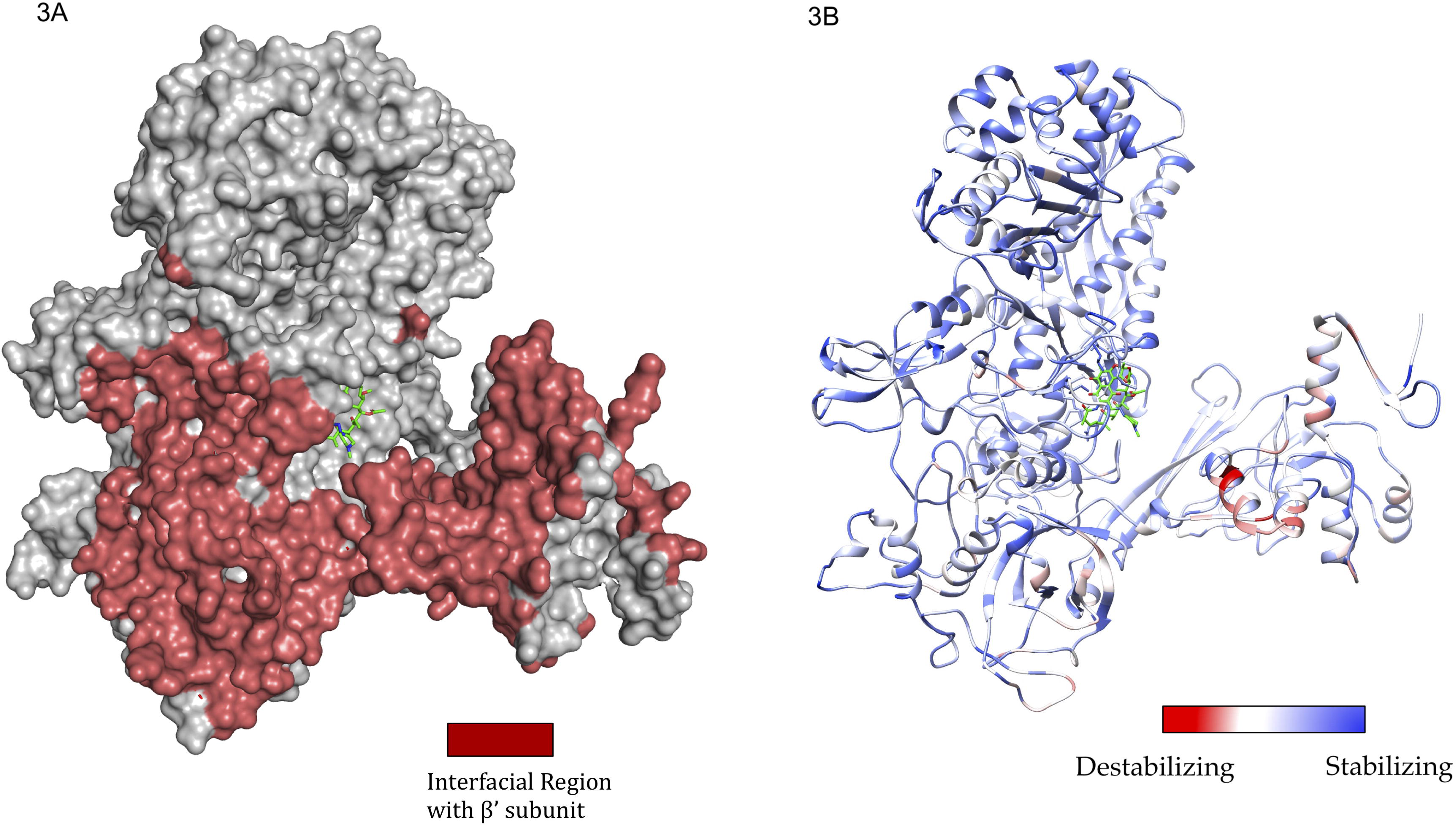
**[A]** The interfacial region of the β subunit of RNAP highlighted in Maroon. [B]. The maximum destabilizing effect a mutation can induce on the interface stability is predicted by mCSM-PPI and mapped on the structure. Red indicates regions that are highly destabilized by mutations (−5.108 Kcal/mol) while the blue indicates stable regions.

### Relative sidechain solvent accessibility (RSA), depth, residue-occluded packing density and protein stability

The difference in relative solvent accessibility between wild type and the mutant residue for all the mutations were calculated using SDM. While analyzing the maximum destabilizing mutations among all the possible mutations at each residue position, it was noted that maximum destabilizing mutants at 751 residue positions (66.79%) show increases in RSA. The maximum destabilizing mutants at rest of the 375 residue positions indicated a decrease in RSA. Among the maximum destabilizing mutants at 751 residue positions which showed an increase in RSA, 551 were hydrophobic and 121 substitutions within 551 were from polar/charged (wildtype) to hydrophobic residues (mutants). As mutant hydrophobic residues with increased solvent accessibility often destabilize the protein [38], the destabilizing effects of these mutations ranged from −1.021 to −4.311 kcal/mol. Additionally, these substitutions resulted in a decrease in residue-depth [28] (ranging from 0.01 Å to 1.83Å), which is concomitant with the increase in solvent accessibility. These changes in RSA and depth at the rifampin-binding site are depicted in Fig 4A & B.

**Fig 4:**
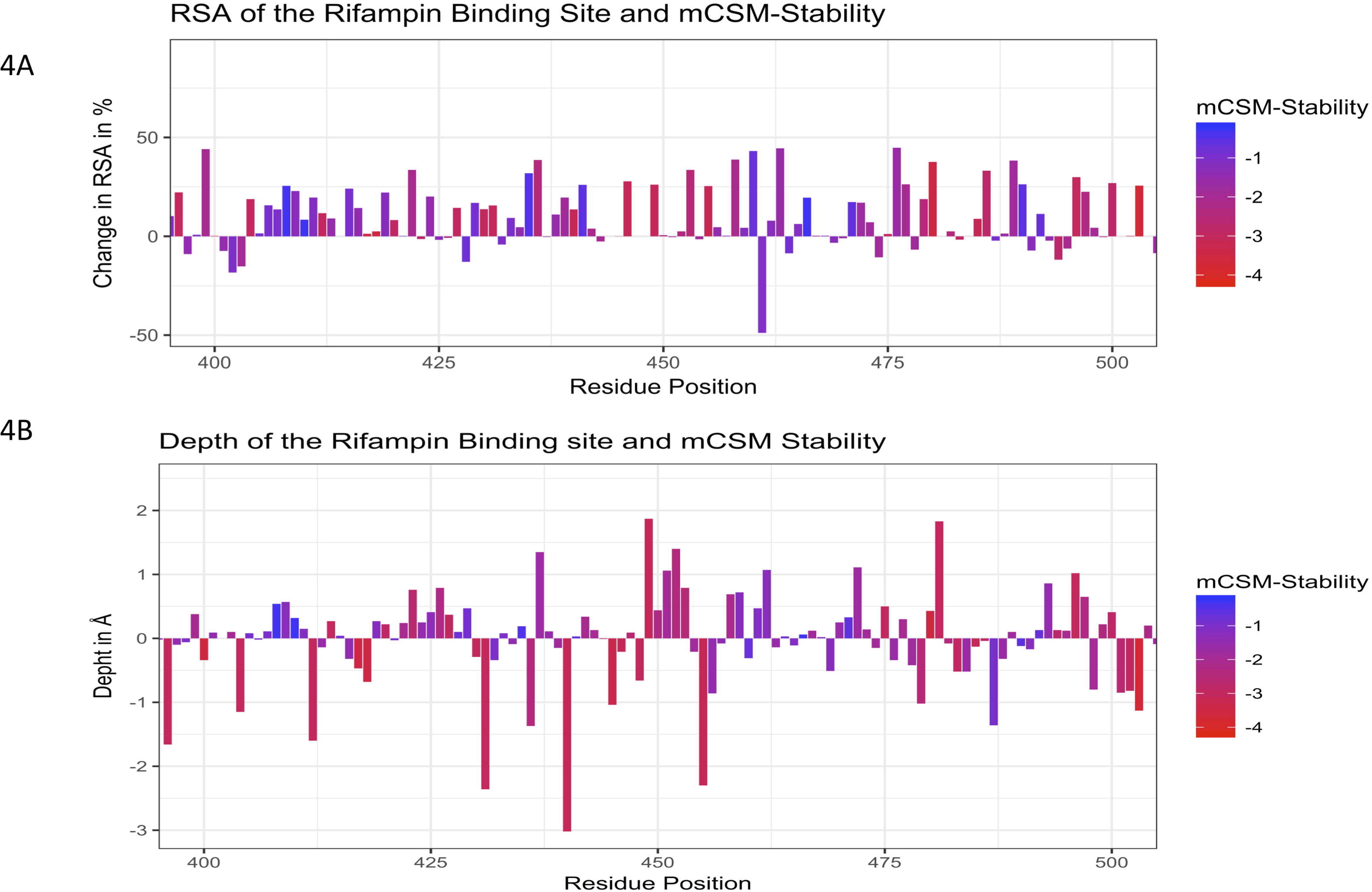
**[A]** Change in relative solvent accessibility for maximum destabilizing mutants in the rifampin binding pocket (mCSM). [B]. Change in depth of the highly destabilizing mutant residue in the rifampin binding pocket (mCSM).

From the maximum destabilizing mutations at all the 1126 positions, mutations at 586 (52.04%) residue positions resulted in increase in depth that ranged from 0.01 to 2.46Å. Mutants were generated using ANDANTE a program that follows χ angle conservation rules to place the sidechains of the mutant residue without any steric clashes. This is followed by energy minimization. Hence the change in depth is attributed to the buriedness of the residue and not just the natural change from a larger to a smaller amino acid. The decrease in depth in the remaining 540 (47.95%) residues ranged from 0.1 to 3.02Å. Similarly, the residue-occluded packing density [28] increased at 539 residue positions (47.86%). These changes in RSA and depth are mapped as attributes on to the structure of the β-subunit of RNAP and it was noted that most of the residues that line the active center cleft have increases in RSA upon mutation. Decrease in depth was noted in residues at the rifampin-binding pocket and at the subunit interfaces (Fig 5A & B).

**Fig 5:**
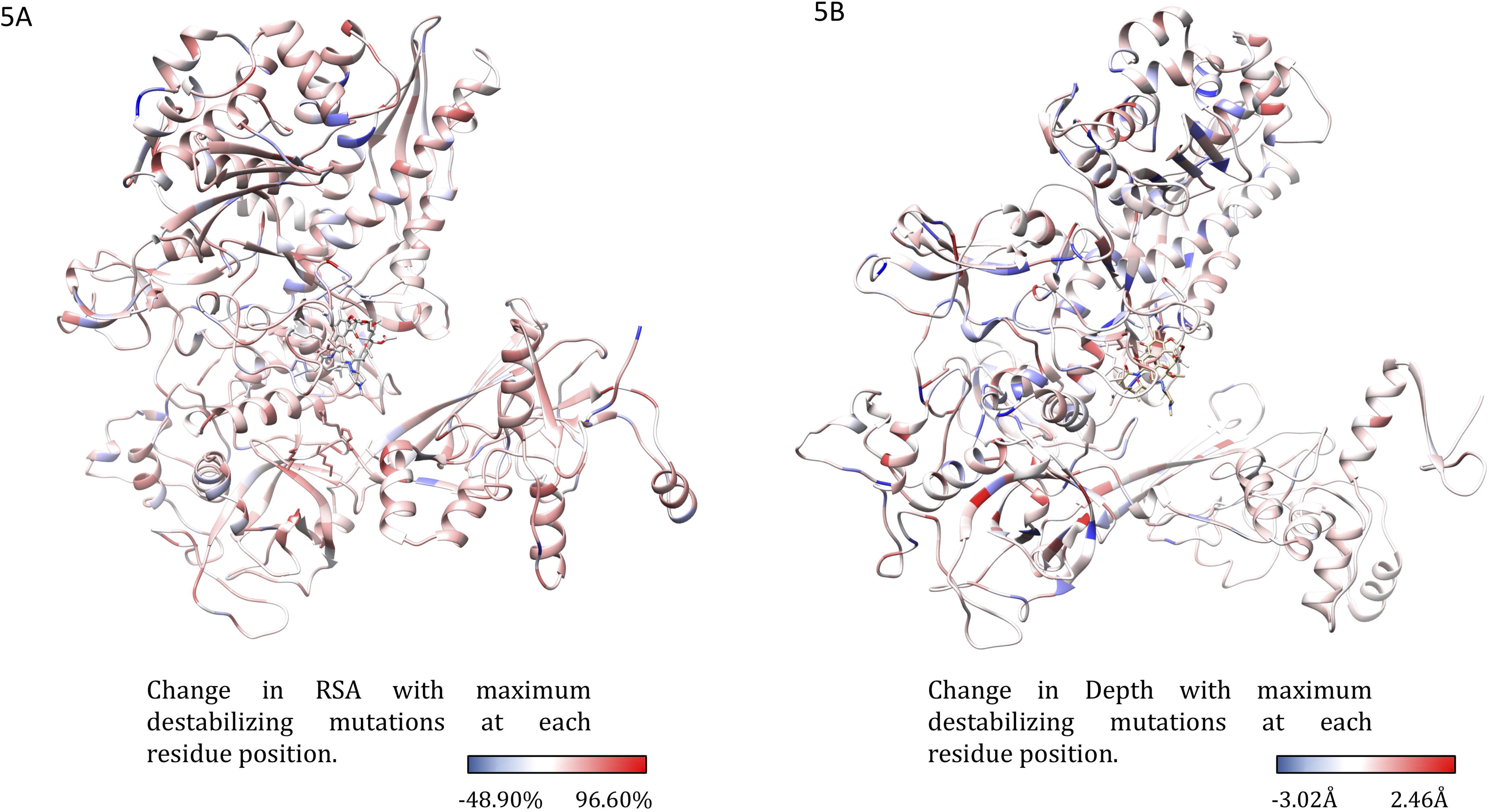
**[A]** The change in relative side chain solvent accessibility with mutations was mapped on to the structure. Blue indicates a decrease in RSA while red indicates an increase. **[B]**The changes in depth with highly destabilizing mutations at each residue position was also mapped on the structure.

### Substitutions to aspartate predominate mutations that destabilize the β subunit-RNA affinity in RNAP

The effects of mutations on β subunit-RNA affinity was estimated using mCSM-NA2. Substitutions to aspartate residues were most common among mutations that highly destabilize β subunit-RNA interactions in RNAP. The mutant aspartate residues induced π-π interactions with the nucleotides in RNA either by stacking or by nucleotide-edge T-shaped and amino-edge T-shaped interactions. Aspartate being an acyclic π-containing amino acid, readily forms nucleotide (edge) amino (edge) or nucleotide (face) and amino-acid (edge) interactions. This ability of acyclic amino acids like arginine, glutamic acid and aspartic acid to form a variety of charged-π interactions with nucleotides in mutants may impact the orientation of RNA molecules in active center cleft of RNAP leading to loss or gain in function. Approximately, 93%of the highly destabilizing mutations at each RNA-interacting residue are substitutions to aspartate. Mutations to glutamate were also noted in 6.83% and additionally one each of methionine, proline and threonine mutations indicated highly destabilizing effects.

### Substitutions to arginine predominate mutations that destabilize β subunit-rifampin affinity

Systematic mutations in the set of 70 residues that lie 10 Å from the rifampin binding site reveal that highly destabilizing mutations are primarily arginine and glutamate substitutions (mCSM-lig). In the binding site R173, R454, R465 and R613 form hydrogen bonds and a network of interatomic interactions with rifampin that stabilize the molecule in the binding site [4], Introduction of additional arginine residues by mutations may influence the stability and orientation of rifampin in the binding site. In predicted mutations S437R and G456R, arginine forms an intricate network of interactions with surrounding aromatic amino acids changing the shape of the binding pocket and leading to a loss in rifampin interactions (rifampin retains only two polar contacts with Q438 and F439 where as wild-type has around five hydrogen bonds). The effects of mutations on RNA and rifampin affinity as predicted by mCSM-NA2 and mCSM-lig were mapped on to the structure (Fig 6A & B).

**Fig 6:**
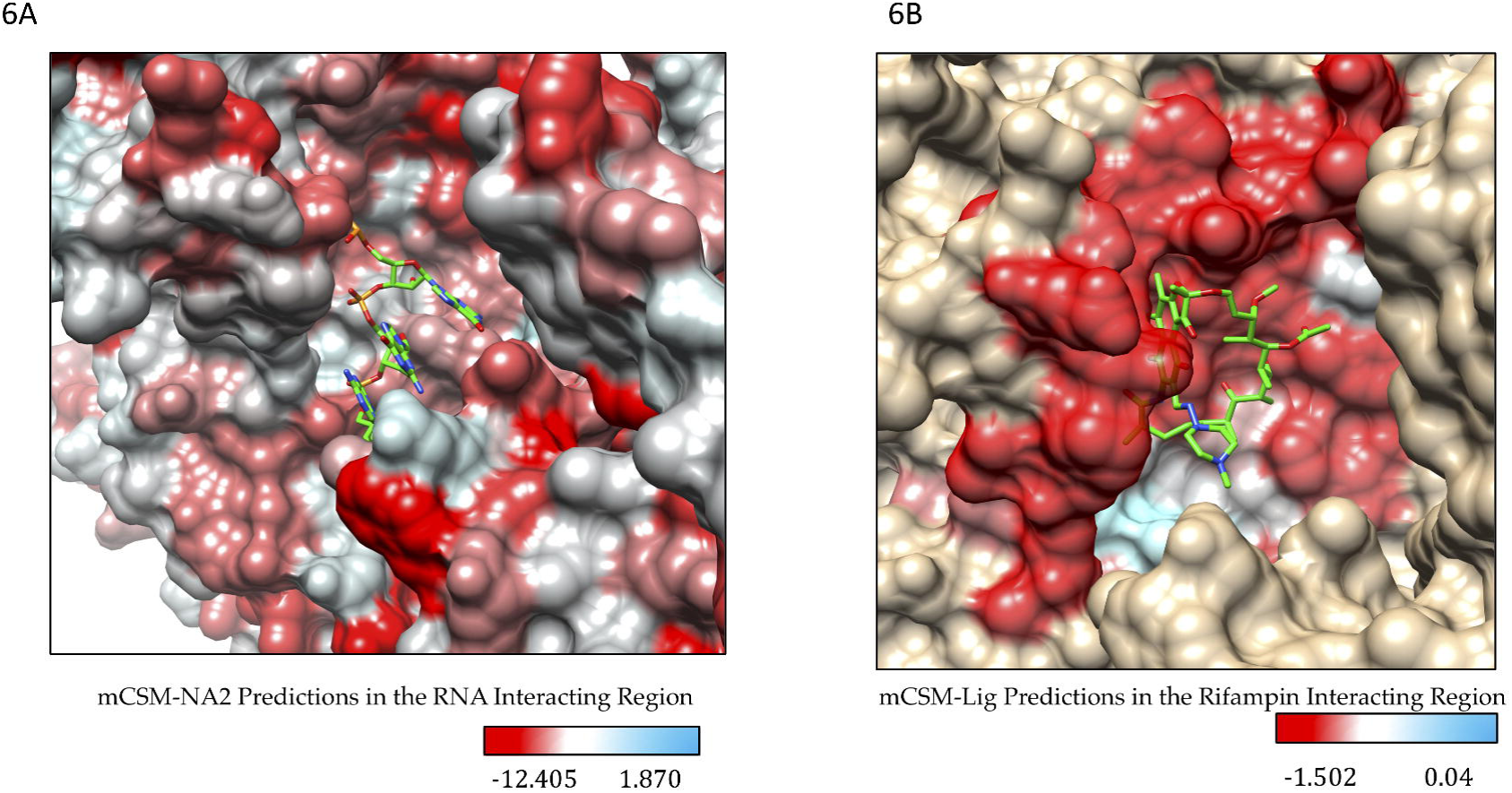
**[A]** Stability changes in β subunit-RNA and β subunit-rifampin **[B]** interactions due to mutations in the binding sites as predicted by mCSM-NA2 and mCSM-lig. The maximum destabilizing effect a mutation can cause at each residue position in the binding site is depicted on the structure.

### Detrimental Mutations

Six residues were chosen based on the following characteristics and the structural effects of systematic mutations at each residue position were analyzed (Table-1)

- Mutations that highly destabilize rifampin binding (at wildtype S437 & G459)
- Experimentally identified and validated mutations that highly destabilize rifampin binding ((at wildtype H451 & P489)[9,10].
- Predicted extremely detrimental mutations for protein stability, and protein-protein and protein-nucleic affinities (at wildtype K884 & H1035).

**Table 1:** Detrimental mutations and their corresponding stability changes that influence holoenzyme assembly, rifampin and RNA interactions.

| Method | Wild-Type Residue | Residue Position | Average Stability Effect | Maximum Stabilizing Effect | Mutant Residue | Maximum Destabilizing Effect | Mutant Residue |
| --- | --- | --- | --- | --- | --- | --- | --- |
| mCSM-Stability | S | 437 | -0.795 | -0.072 | L | -1.701 | H |
|  | H | 451 | -1.214 | -0.104 | Y | -1.898 | S |
|  | G | 459 | -0.713 | -0.381 | V | -1.201 | W |
|  | P | 489 | -1.135 | -0.507 | R | -1.771 | G |
|  | K | 884 | -1.227 | -0.190 | L | -2.298 | S |
|  | H | 1035 | -0.419 | 0.600 | Y | -1.421 | G |
| mCSM-PPI | S | 437 | -0.254 | 0.395 | H | -0.820 | R |
|  | H | 451 | -0.652 | -0.050 | S | -1.451 | M |
|  | G | 459 | -0.397 | 0.237 | H | -1.042 | R |
|  | P | 489 | -0.738 | -0.138 | W | -1.372 | R |
|  | K | 884 | -0.105 | 0.160 | D | -0.685 | R |
|  | H | 1035 | -0.754 | 0.115 | W | -1.726 | R |
| mCSM-NA | S | 437 | -1.538 | 4.922 | W | -3.857 | D |
|  | H | 451 | -1.300 | 5.147 | W | -3.632 | D |
|  | G | 459 | 2.289 | 8.556 | W | -0.221 | D |
|  | P | 489 | 1.926 | 8.195 | W | -0.582 | D |
|  | K | 884 | 0.221 | 6.647 | W | -2.130 | D |
|  | H | 1035 | 0.847 | 7.295 | W | -1.484 | D |
| mCSM-Lig | S | 437 | -0.646 | -0.484 | L | -1.062 | R |
|  | H | 451 | -0.510 | -0.076 | W | -0.777 | E |
|  | G | 459 | -0.981 | -0.715 | A | -1.236 | R |
|  | P | 489 | -0.598 | -0.254 | L | -0.917 | R |
|  | K | 884 | -0.156 | -0.368 | D | -0.925 | R |
|  | H | 1035 | -0.121 | 0.097 | V | -0.501 | E |
| SDM | S | 437 | 0.087 | 2.320 | V | -1.900 | P |
|  | H | 451 | -0.756 | 1.290 | L | -2.800 | G |
|  | G | 459 | -2.842 | -1.780 | V | -3.800 | P |
|  | P | 489 | -0.432 | 1.440 | Y | -1.070 | E |
|  | K | 884 | 0.108 | 1.270 | V | -1.820 | P |
|  | H | 1035 | -0.200 | 0.590 | V | -1.410 | P |
| MAESTRO | S | 437 | -0.21 | -0.14 | K | 0.24 | F |
|  | H | 451 | -0.12 | -0.05 | G | 0.22 | R |
|  | G | 459 | -0.23 | -0.17 | S | 0.33 | W |
|  | P | 489 | -0.26 | -0.22 | H | 0.31 | M |
|  | K | 884 | -0.20 | -0.14 | G | 0.25 | M |
|  | H | 1035 | -0.27 | -0.25 | P | 0.31 | Y |
| CUPSAT | S | 437 | 2.70 | 7.98 | I | -1.12 | G |
|  | H | 451 | 2.01 | 6.92 | W | -3.25 | K |
|  | G | 459 | -2.51 | 5.00 | K | -5.53 | C |
|  | P | 489 | -2.76 | -0.84 | A | -5.47 | M |
|  | K | 884 | -2.99 | 3.42 | I | -8.03 | H |
|  | H | 1035 | -1.07 | 2.15 | C | -3.23 | Y |
| Imutant 2.0 Structure | S | 437 | 4.05 | 9.00 | A | 1.00 | F |
|  | H | 451 | 6.00 | 8.00 | G | 3.00 | L |
|  | G | 459 | 6.63 | 9.00 | N | 3.00 | I |
|  | P | 489 | 7.11 | 9.00 | G | 3.00 | L |
|  | K | 884 | 6.42 | 9.00 | G | 2.00 | M |
|  | H | 1035 | 4.63 | 8.00 | G | 2.00 | L |
| PROVEAN | S | 437 | -4.79 | -3.00 | A | -7.00 | W |
|  | H | 451 | -8.66 | -5.73 | Y | -10.37 | C |
|  | G | 459 | -8.10 | -6.00 | A | -10.00 | L |
|  | P | 489 | -9.04 | -7.99 | A | -10.99 | F |
|  | K | 884 | -5.97 | -2.91 | R | -7.75 | C |
|  | H | 1035 | -8.98 | -5.79 | Y | -10.61 | C |
| Imutant 2.0 Sequence | S | 437 | 4.47 | 7.00 | F | 0.00 | H |
|  | H | 451 | 3.21 | 7.00 | P | 0.00 | F |
|  | G | 459 | 3.53 | 7.00 | H | 0.00 | A |
|  | P | 489 | 6.89 | 9.00 | G | 5.00 | L |
|  | K | 884 | 3.53 | 8.00 | V | 0.00 | G |
|  | H | 1035 | 2.95 | 6.00 | G | 0.00 | V |
| FOLDX4 | S | 437 | 2.79 | -1.44 | I | 12.39 | R |
|  | H | 451 | 1.78 | -0.74 | L | 4.39 | W |
|  | G | 459 | 9.14 | 3.96 | A | 20.76 | H |
|  | P | 489 | 3.04 | 2.11 | N | 4.79 | R |
|  | K | 884 | 1.06 | -2.12 | Y | 9.77 | L |
|  | H | 1035 | 0.77 | -1.47 | P | 5.69 | Y |
| ENCoM | S | 437 | -0.44 | 0.48 | G | -1.50 | W |
|  | H | 451 | 0.34 | 0.97 | G | -0.46 | W |
|  | G | 459 | -0.91 | -0.29 | A | -1.55 | W |
|  | P | 489 | -0.16 | 0.14 | G | -0.82 | F |
|  | K | 884 | 0.18 | 0.96 | G | -0.60 | W |
|  | H | 1035 | 0.19 | 0.73 | G | -0.26 | W |
| DynaMut | S | 437 | 2.87 | 6.99 | L | -2.08 | G |
|  | H | 451 | -0.74 | 2.17 | Y | -3.43 | T |
|  | G | 459 | 1.93 | 3.29 | N | -0.25 | S |
|  | P | 489 | 0.94 | 3.26 | F | -0.72 | S |
|  | K | 884 | 0.14 | 3.69 | W | -1.87 | E |
|  | H | 1035 | 0.21 | 2.38 | W | -2.29 | G |

### Detrimental mutations in the rifampin binding site

We have noted that any mutation at rifampin-interacting residues S437, H451, R454, S456, L458, G459, R465, P489, P492 and N493 destabilize protein ligand affinity (mCSM-lig). Of these we have chosen wild-type residues H451 and P489, which are experimentally identified mutations, and wild-type residues S437 and G459, which are computationally predicted (only one mutation was experimentally identified at residue position S437L as reported by us earlier [4], and this has destabilizing effects on the overall stability and affinity to rifampin).

### S437

Serine at position 437 in the wild-type structure forms mainchain and sidechain hydrogen bond interactions with S434, G432 and R173. The residue has a network of proximal polar interactions and hence stabilizes the rifampin-binding pocket. It was noted that any mutation at this position reduces rifampin affinity (mCSM-lig) and stability of the β subunit (mCSM) (Supplementary-Table 1) (Fig 7A). The maximum destabilizing effect was noted for substitution to histidine (−1.701 kcal/mol (mCSM)) where it forms hydrogen bond interactions with S434 and Q438, aromatic interactions with F431, and a network of ring-ring and π interactions with the surrounding residues which might largely effect the shape of the binding pocket (Fig 7B). Substitution with leucine causes a minimal destabilizing effect (−0.072 kcal/mol(mCSM)) and stability effects of all the other amino acid substitutions range from −0.072 to −1.701 kcal/mol (mCSM).

**Fig 7:**
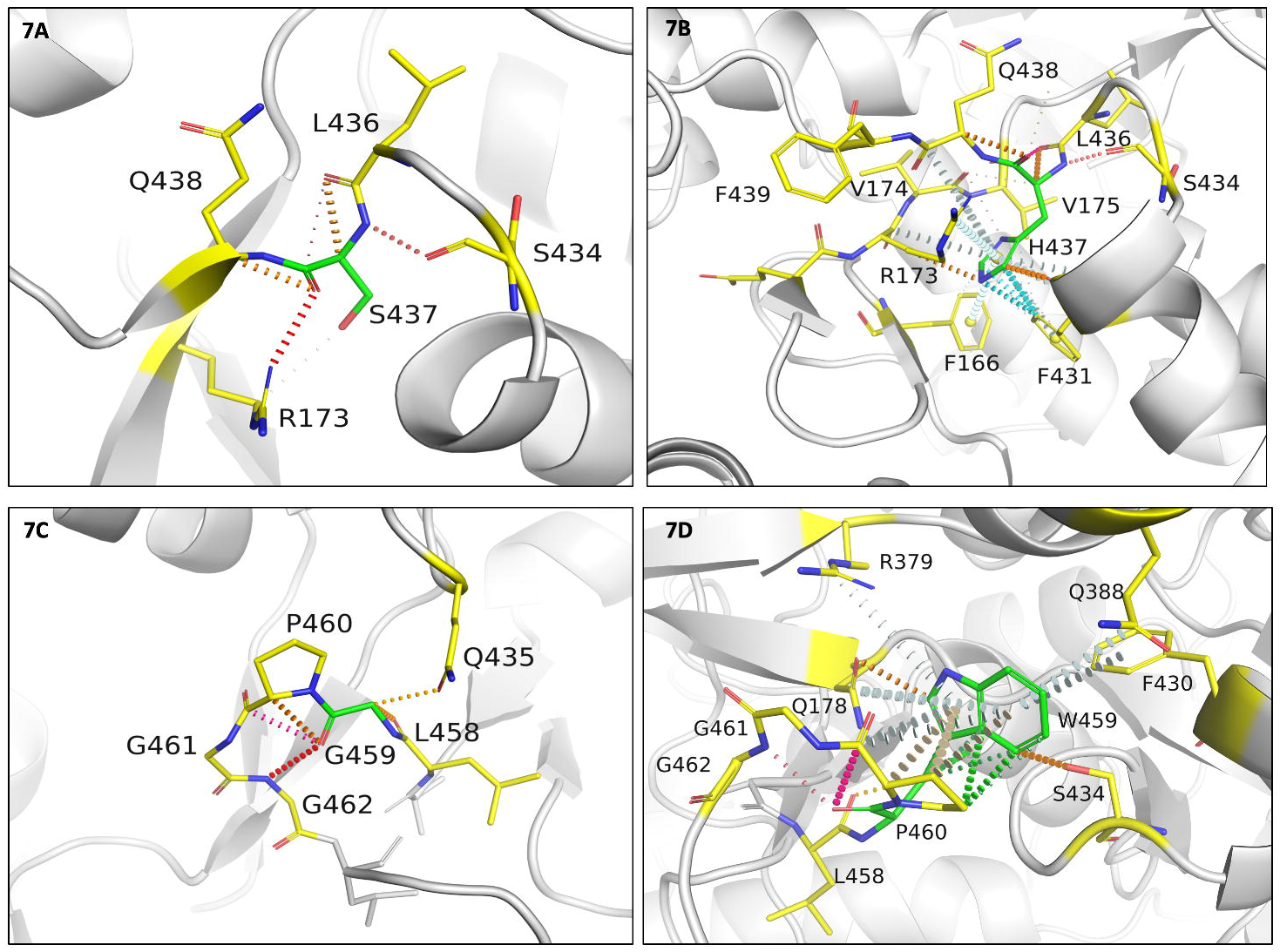
**[A]** Interactions of S437 with the surrounding residue environment in the wildtype and of H437 in the S437H mutant **[B]**. **[C]** Interactions of G459 with the surrounding residue environment and **[D]** W459 in the mutant G459W. The red dotted lines represent hydrogen bonds. Orange dotted lines represent weak hydrogen bond interactions. Ring-Ring and intergroup interactions are depicted in cyan. Aromatic interactions are represented in sky-blue and carbonyl interactions in pink dotted lines. Green dotted lines represent hydrophobic interactions.

S437 is located at 3.3 Å from the interface of β and β’ subunits. Arginine substitution destabilized the interface with the predicted interface stability change of −0.820 kcal/mol (mCSM-ppi). In the wild-type structure, S437 is located 11.9 Å from the closest nucleic acid molecule but is present on the helix that interacts with both DNA and transcribing RNA in the active center cleft. An aspartate substitution destabilizes the protein-RNA interaction with predicted affinity change of-3.857 kcal/mol (mCSM-NA2). S437 is located 4.0 A from rifampin and forms proximal interactions with rifampin. However, S437 forms hydrogen bond interactions with S434 and R173 that are important for the attachment of rifampin to the binding pocket. The S437R mutation disrupts the hydrogen bond interactions with S434 and R173 which in-turn impact stability of rifampin in the binding pocket (−1.062 kcal/mol (mCSM-lig)).

### G459

Glycine at position 459 forms hydrogen bonds with Q435, L458 and G462, and carbonyl interactions with the P460. G459 is present 4.6 Å away from rifampin and is involved in hydrogen bonds with residues that interact with rifampin (Fig 7C). A tryptophan substitution largely destabilizes the binding pocket by the incorporation of hydrophobic and π interactions with the surrounding residues. It forms side-chain hydrophobic interactions with L436, L384 and F430. It also forms a ring–ring interaction with F430, an atom-ring interaction with L384 and intergroup interactions with Q178 and Q388. It forms multiple hydrogen bonds with the surrounding residues, which may impact the orientation of the binding pocket and destabilize the protein (Fig 7D).

### Experimental Mutations that highly destabilize rifampin binding

From the 40 mutations that are reported from different rifampin-resistant leprosy clinical isolates, we have chosen two residues where mutations are extremely detrimental to protein stability, protein ligand affinity, protein nucleic affinity and protein subunit interfaces. Substitutions at H451 and P489 were studied in detail.

### H451

H451 in the wild-type structure lies 3.7Å from rifampin and 4.1Å from the interface. This residue forms cation -π interactions with guanidinium group of R454, which in turn forms polar interactions with rifampin (Fig 8A). Additionally, H451 makes two hydrogen bonds with mainchain amino group of R454 and oxygen atom of S447. Mutations at this residue site largely impact the stability and ligand binding. Substitution to serine induced a change in stability of the protein with a decrease in Gibbs free energy of −1.898 kcal/mol and a network of π interactions that are present in the native structure were lost in the mutant (Fig 8B).

**Fig 8:**
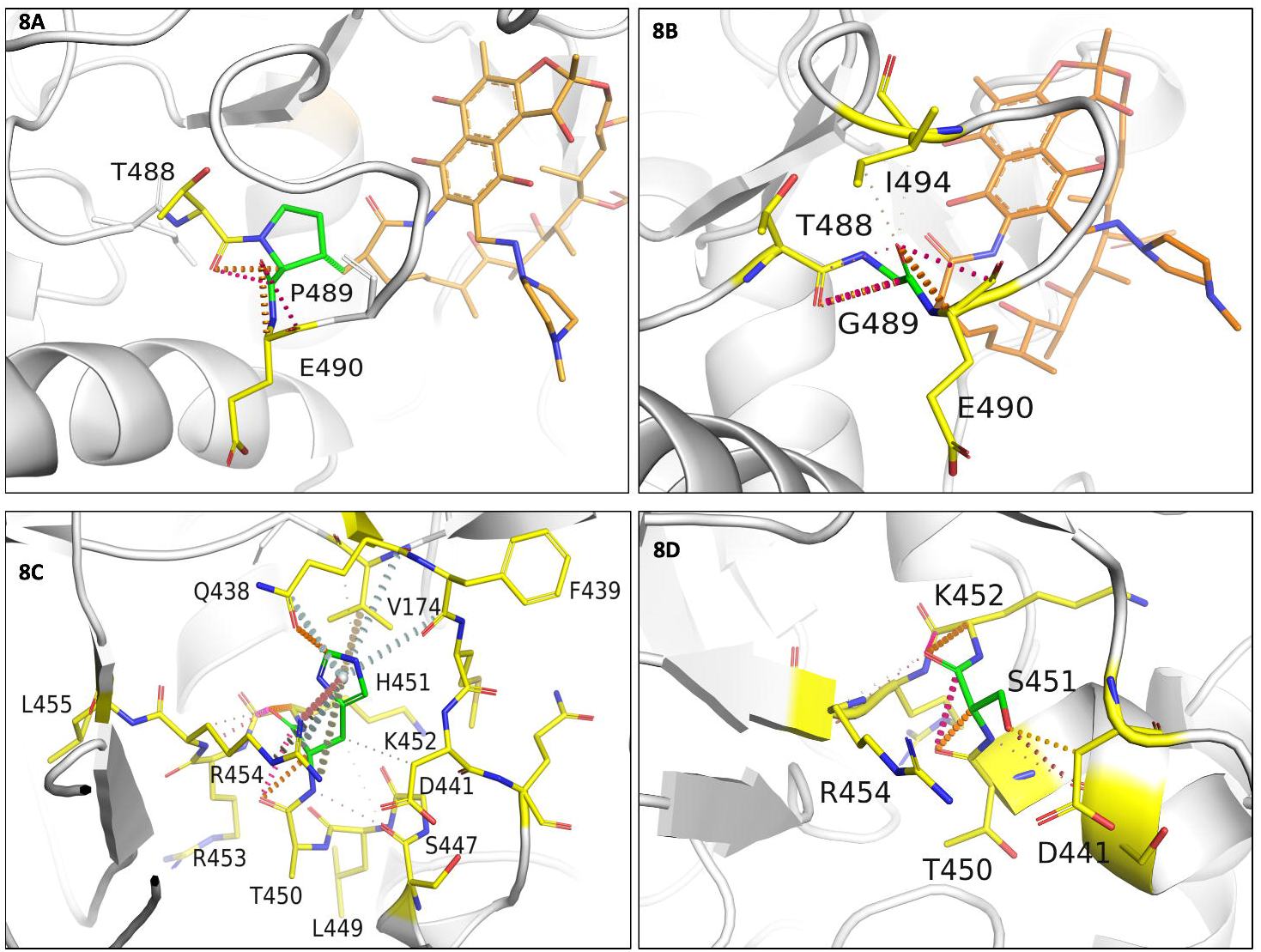
**[A]** Interactions of P489 with the surrounding residue environment in the wildtype and of G489 in the P489G mutant **[B]. [C]** Interactions of H451 with the surrounding residue environment and **[D]** S451 in the mutant H451S.

Methionine substitution destabilizes β - β’ subunit interface and leads to a change in free energy of −1.451 Kcal/mol. Methionine forms carbonyl interactions with K452 and T450, a hydrophobic interaction with Q438 and weak hydrogen bond interactions with rifampin. Although histidine or methionine do not directly interact with the residues of the β’ subunit, the changes in the network of π-interactions coupled with the addition of hydrophobic interactions with proximal residues in the interface may change their binding patterns leading to destabilization of the interface.

Substitution with glutamic acid induces a destabilizing effect on the β subunit-rifampin interaction. E451 forms weak hydrogen bond, carbonyl and proximal hydrophobic interaction but does not form any bonds with rifampin, unlike the wild-type residue that forms proximal hydrogen bonds with rifampin.

### P489

Proline at position 489 is present in a loop which is in close proximity to rifampin and forms hydrophobic interaction with rifampin and weak hydrogen bond interactions with T488 and Q490 (Fig 8C). Mutations at the position 489 were reported in rifampin-resistant leprosy patients from Thailand [9]. Glycine substitution destabilizes the protein (−1.771kcal/mol) leading to a loss of hydrophobic interaction with rifampin. Weak hydrogen bond and carbonyl interactions, however, were retained in the mutant model (Fig 8D). Arginine substitution destabilizes interface and rifampin affinities, with predicted stability changes of −1.372 and −0.917 kcal/mol respectively. FoldX predicted a large change in stability of 4.79 kcal/mol for difference between mutant and wild types, which is highly destabilizing. FoldX optimizes the sidechains and moves the structure to a lowest energy state (usually represented as a negative value) and hence the difference between two negative energy values of wild and mutant is destabilizing.

### Extremely Detrimental Mutations

Mutations at residues positions K884 and H1035 were considered to be extremely detrimental. These residues lie in close proximity to the interface, nucleic acids and rifampin. Substitutions at these sites destabilize protomer, protein-protein interfaces (both the residues reside at the subunit interface), protein-nucleic acid and protein-ligand affinities. Both empirical (FoldX) and knowledge based (mCSM and SDM) methods predicted destabilizing effects.

### K884

K884 is located 3.2 Å from the interface, 3.3 Å from the nucleic acid and 8.6Å from rifampin. Lysine forms mainchain hydrogen bonds with L1033 and proximal hydrophobic interactions with H1035 and V894. It also forms a cation -π interactions with H1035 and most importantly a sidechain proximal hydrogen bond with the sugar phosphate group of guanine (second) nucleotide in the RNA transcript. This interaction is critical for maintaining the RNA interaction with rifampin in order to induce steric clash on the adjacent nucleotide and halt transcription (Fig 9A). Serine substitution at this site results in the loss of these vital interactions. S884 forms weak van der Waals interactions with D883 and L885 and hydrogen bonds with L1033 and H1035. Interactions with RNA backbone are lost in the mutant (Fig 9B). The mutant is destabilized with a predicted stability change of −2.298 kcal/mol.

**Fig 9:**
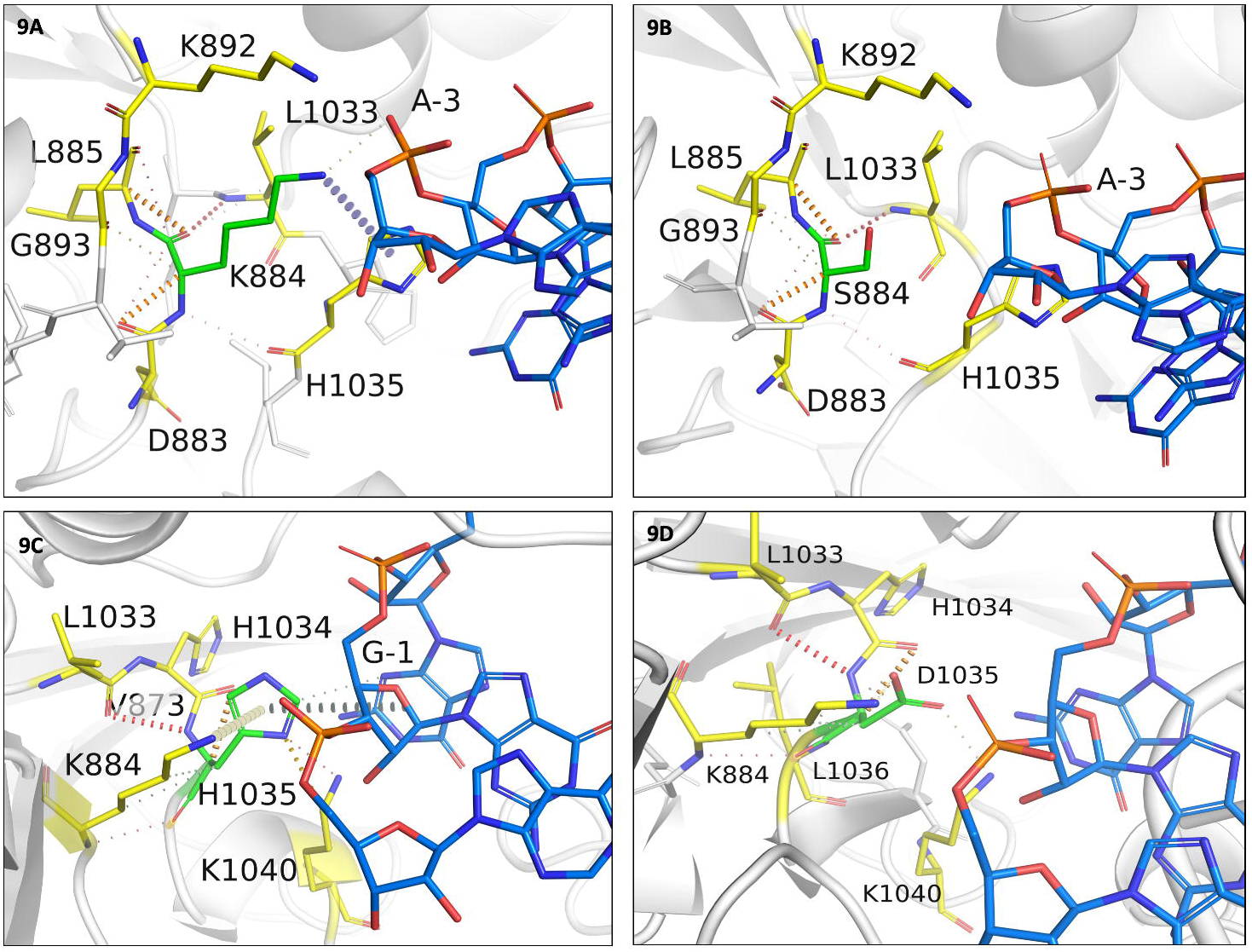
**[A]** Interactions of K884 with the surrounding residue environment in the wildtype and of S884 in the K884S mutant **[B]. [C]**Interactions of H1035 with the surrounding residue environment and **[D]** D1035 in the mutant H1035D. The blue dotted lines represent cation-π interaction.

Aspartate substitution at this site destabilizes RNA affinity with a change of −2.130 kcal/mol and the mutant residue forms hydrogen bonds with L1033 and H1035, and hydrophobic interactions with V894.

### H1035

Histidine at position 1035 is located 3.5 Å from the interface and RNA, and 8.8Å away from rifampin. It forms a network of π interactions with the surrounding residues. The ring-ring π interactions with the fused pyrimidine-imidazole ring of guanine in the first nucleotide of RNA transcript is vital to the orientation of RNA transcript in the active center cleft (Fig 9C). These interactions are lost in mutations especially with non-aromatic amino acids. It was also noted that aspartate substitution largely destabilizes β subunit-rifampin affinity (Fig 9D).

### Impact of Mutations on Flexible conformations

The stability changes between the wildtype and each mutant in lowest energy conformation were calculated by FoldX and have a Pearson’s correlation coefficient (“r” value) of 0.38 with other predictors mCSM and SDM. Although FoldX does not probe backbone conformational changes, it optimizes the sidechain rotamers of the mutant residues to attain a low energy state and calculates the change in free energy between the states. We further sampled the fully flexible conformers of the β subunit and estimated changes in vibrational entropy ΔS and protein stability using ENCoM. A linear combination of vibrational entropy ΔS by ENCoM and enthalpy changes by FoldX was used to calculate stability changes. ENCoM predicted highly destabilizing mutations in the rifampin binding and RNA interacting sites in the active center cleft of the holoenzyme. DynaMut predictions correlated with ENCoM values at an r value of 0.56. The average change in stability predicted by ENCoM and DynaMut for any mutation at each residue in the β subunit was mapped on the structure (Fig 10A and B).

**Fig 10:**
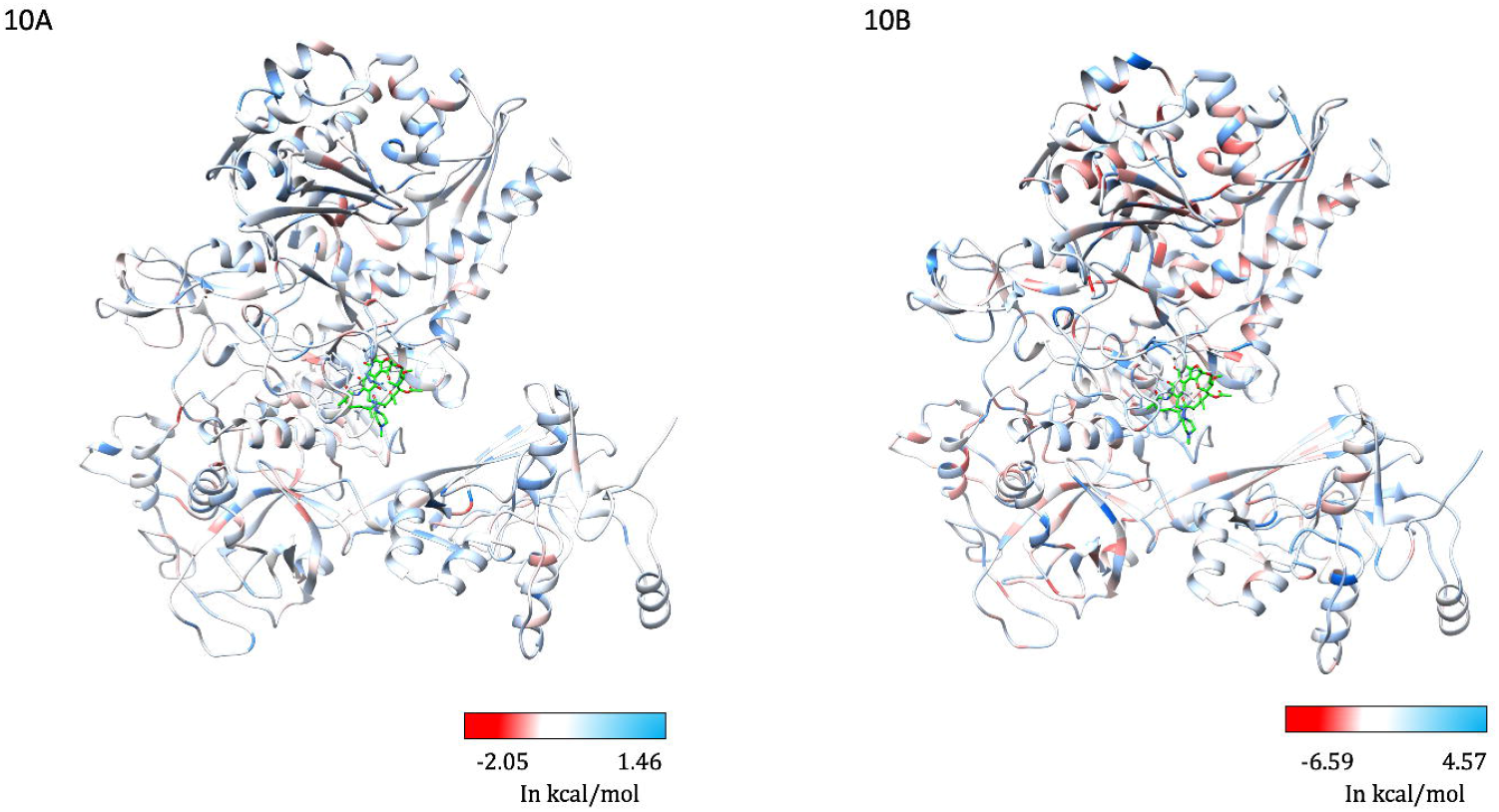
**[A]** The maximum destabilizing effects on the protein stability, a mutation can induce at each residue position in the flexible conformations (as predicted by ENCoM [A] and DynaMut [B], are mapped on the structure. Regions in red represent highly destabilizing while the blue regions are relatively stable with mutations.

### Stability changes and fragment hotspot maps

Hotspots were mapped on the structure and colored with maximum destabilizing effects caused by any mutations at each residue site. The regions of the β subunit that are least impacted by mutations (mutation coolspots) are overlaid with fragment hotspots. The site B (Fig 11), which is in close proximity to the RNA binding region and is a pocket at the β-β’ subunit interface, is least impacted by mutations and has a hotspot at the contouring score of 17 with donor, apolar and acceptor regions [21]. Secondly, the site A, although located away from the catalytic core of the enzyme, is present in the path of entry/exit point for template DNA into the holoenzyme complex and a small molecule interaction at this site can potentially impact template DNA interactions or induce conformational change in the crab-claw-shaped β subunit leading to disruption in the holoenzyme assembly.

**Fig 11:**
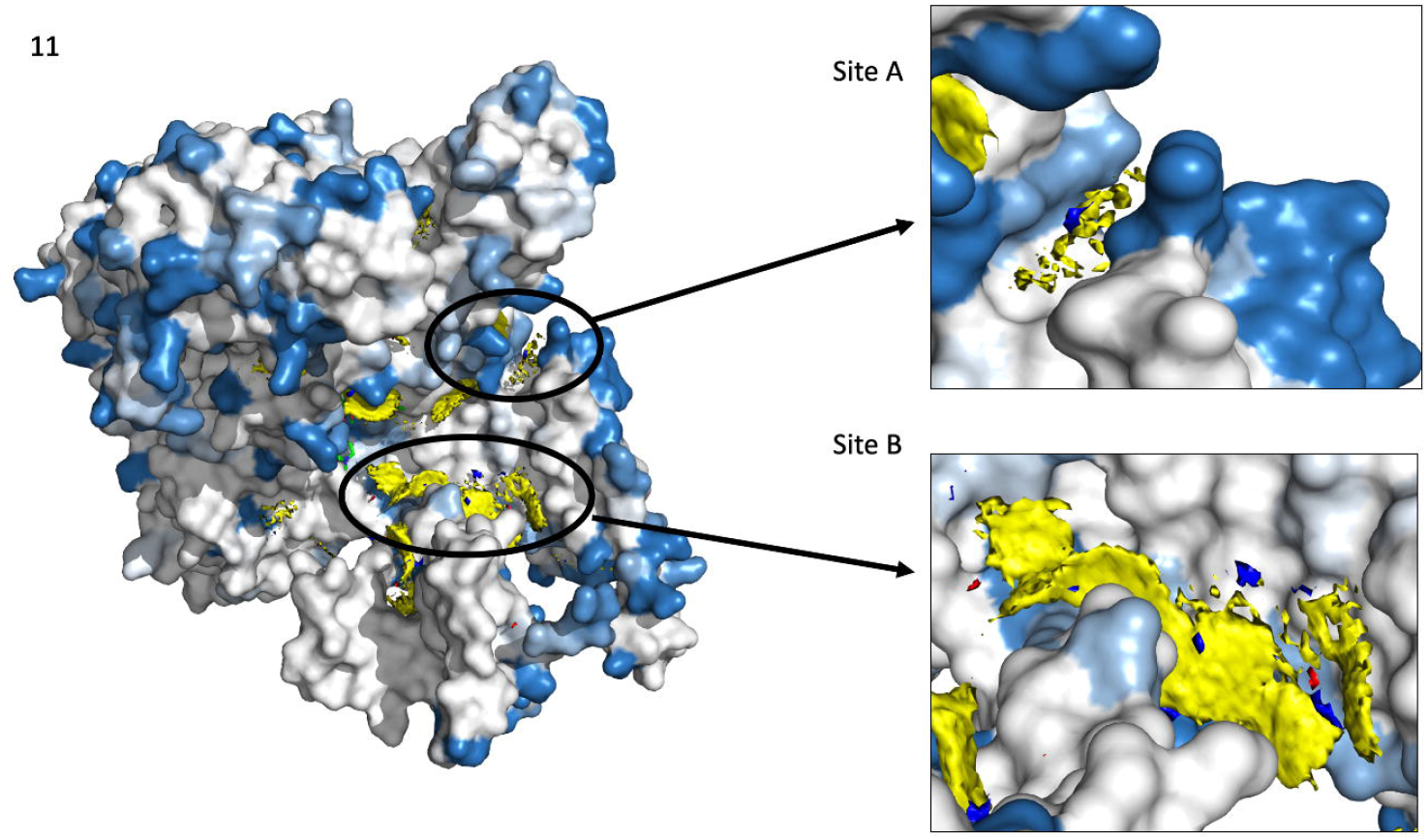
Fragment hotspots were mapped on the structure which was coloured with maximum destabilizing effects of systematic mutations at each residue positions. Blue represents regions which are least impacted by any mutations. Stable and potential small molecule binding sites “A” and “B” are depicted on the structure.

## DISCUSSION

In the absence of a rapid and an effective laboratory-based diagnostic tool for determining drug resistance in leprosy, identification of mutations known to confer resistance to individual drugs in multidrug therapy remains an appropriate approach for diagnosing drug resistance. Associations between mutations in drug targets and clinical resistance to individual drugs in MDT are often validated by mouse-footpad experiments in which, resistant strains (with known mutations) are propagated in the hind footpads of mice (cross-bred albino) in the presence of drugs under study [4]. Rifampin resistance is widespread in tuberculosis with annual global incidence of ~600,000 cases and since the molecular mechanisms of drug action are similar in both tuberculosis and leprosy, it is expected that rifampin-resistant strains of *M. leprae* may also exist in numbers much higher than those reported through various epidemiological studies [10]. Owing to high percentage identity of the β subunit of RNAP of *M. leprae* with that of *M. tuberculosis*, identical mutations that are experimentally proven to confer rifampin resistance in tuberculosis, are considered as likely drug-resistant mutations in leprosy. The experimentally known mutations in *M. leprae* were those identified by DNA sequencing of *rpoB* gene (derived from skin tissues DNA of relapsed/drug resistant leprosy patients) and published in different studies (references for each mutation are listed in Supplementary Table-2). Most of these were validated in either mouse foot-pad experiments or by using surrogate genetic hosts. Any new mutations that emerge will need experimental validation using mouse footpad /other experimental methods, which are time consuming, posing the need for effective alternative solutions to decipher the possible impacts of the mutations on drug-resistance outcomes [39].

Around 40 different rifampin-resistance mutations were noted in *M. leprae* from clinical isolates around the world using amplicon sequencing of rifampin resistance determining region(RRDR)[10]. All of these mutations decrease the stability of rifampin binding to the β subunit of RNAP (Supplementary Table-1) and the mutant strains exhibited normal grown patterns in the mouse footpads when administered with rifampin in doses equivalent to WHO regimen of MB MDT [40]. This indicates that mutations structurally and functionally impact rifampin interactions and the concomitant resistance.

Thermodynamic stability of the proteins essentially influences their function and is largely dependent on the sequence. Missense mutations that lead to amino acid substitutions often impact protein stability, shifting it towards either a stabilized or a destabilized state [7]. Experimental measurements of stability changes in proteins are often challenging especially with large and complex protein machineries like RNAP. However, mutations within each subunit of the RNAP complex, and primarily the rifampin binding β subunit, have clinical implications and influence rifampin-resistance outcomes in mycobacterial diseases[41]. The performance of various structural, sequence and NMA based predictors for predicting protein stability changes upon mutations vary largely in terms of their accuracy and bias[42], but offer a quick and a helpful alternative to understanding the association between mutations and resistance phenotypes[6].

Given the absence of a rapid and experimentally validated system to read the impact of mutations in the β-subunit of RNAP in *M. leprae* with clinical rifampin resistance outcomes in leprosy, we conducted computational saturation mutagenesis to determine regions on the βsubunit that impact the overall stability, protein-subunit interfaces, protein-nucleic and protein-ligand affinities. Being a part of the complex transcriptional machinery in the mycobacterial cell, the compositional and conformational stability of the β-subunit is crucial to binding of DNA template and synthesis of complementary RNA transcript in the active center cleft of the holoenzyme[43,44]. As rifampin blocks the growing RNA transcript through steric occlusion, its binding and orientation in the binding pocket are vital to its function [43]. Mutations within the RRDR impact rifampin interactions and overall stability of the subunit. As noted from Table 1, all the experimentally identified *rpoB* gene mutations from *M. leprae* indicated a destabilizing effect on the protein-ligand affinity. Owing to the robustness of these predictions, we employed an *in-silico* saturation mutagenesis model to understand the impacts of systematic mutations at each residue site of the subunit.

The destabilizing mutations are given preference over mutations that are silent or have minimal effects on the stability. This is to explore and understand the possible structural and functional implications of emerging detrimental mutations (reported or new) that can influence rifampin resistance outcomes in leprosy. We used different structural, sequence and NMA based tools to identify and compare the predictions. mCSM stability predictions had better correlations with the other predictors (SDM (r=0.55), MAESTRO (r=0.61), Imutant 2.0 Structure (r=0.72), CUPSAT (r=0.43), Imutant 2.0 Sequence (r=0.62) and Dynamut (r = 0.61)).

Protocols (Computational Saturation Mutagenesis (CoSM))[45] that use molecular dynamic equilibration, sidechain flips and energy minimization to improve side conformations in mutants enable prediction of stability changes with better accuracy and correlation with the experimentally deciphered stability changes (r=0.9). However, these protocols are computationally intensive and require high performance computing systems and time. CoSM had a similar performance to FoldX, which was used in the current study. Given the large sample, size molecular dynamic equilibration of sidechain rotamers is beyond the scope of this study.

In conclusion, we have deciphered the predicted effects of all possible mutations in the β subunit of RNAP of *M. leprae* using computational saturation mutagenesis model, probing structural, sequence driven and dynamic changes that impact overall stability of the protein, RNA and rifampin affinities. The predicted impacts were mapped onto the structures and highly detrimental mutations are further analyzed for their changes in interatomic interactions. Due to the lack of adequate experimental data on stability changes in β subunit of RNAP upon mutations, we have limited information on the accuracy of the predictions, however, all the prediction tools used in the study are well tested and validated software which are proven to perform with reasonable accuracy and minimal bias on various relevant mutational datasets [34]. To date there were no studies describing the phenotypic resistance/susceptibility outcomes in strains with compensatory mutations in RNAP. Further studies on saturation mutagenesis of the entire RNAP holoenzyme complex may provide comprehensive information on the effects of co-evolving and compensatory mutations in other subunits on rifampin binding and function.

## ACKNOWLEDGEMENTS

Authors would like to thank the rest of the computational biology team at Department of Biochemistry, University of Cambridge for their overarching support and guidance in the data collection and analysis. SCV was supported by American Leprosy Missions Grant (G88726), MJS was supported by a grant from Foundation Botnar working to support children with cystic fibrosis (Project 6063), CHMR and SP were supported by Australian Government Research Training Program Scholarships, DBA was funded by a Newton Fund RCUK-CONFAP Grant awarded by The Medical Research Council and Fundação de Amparo à Pesquisa do Estado de Minas Gerais (MR/M026302/1) and by the Wellcome Trust Programme Grant (200814/Z/16/Z) and supported in part by the Victorian Government’s OIS Program. TLB was supported by the Wellcome Trust Programme Grant (200814/Z/16/Z) and SM was supported by the MRC DBT Grant (RG78439).

---

**Supplementary Figure 1:**
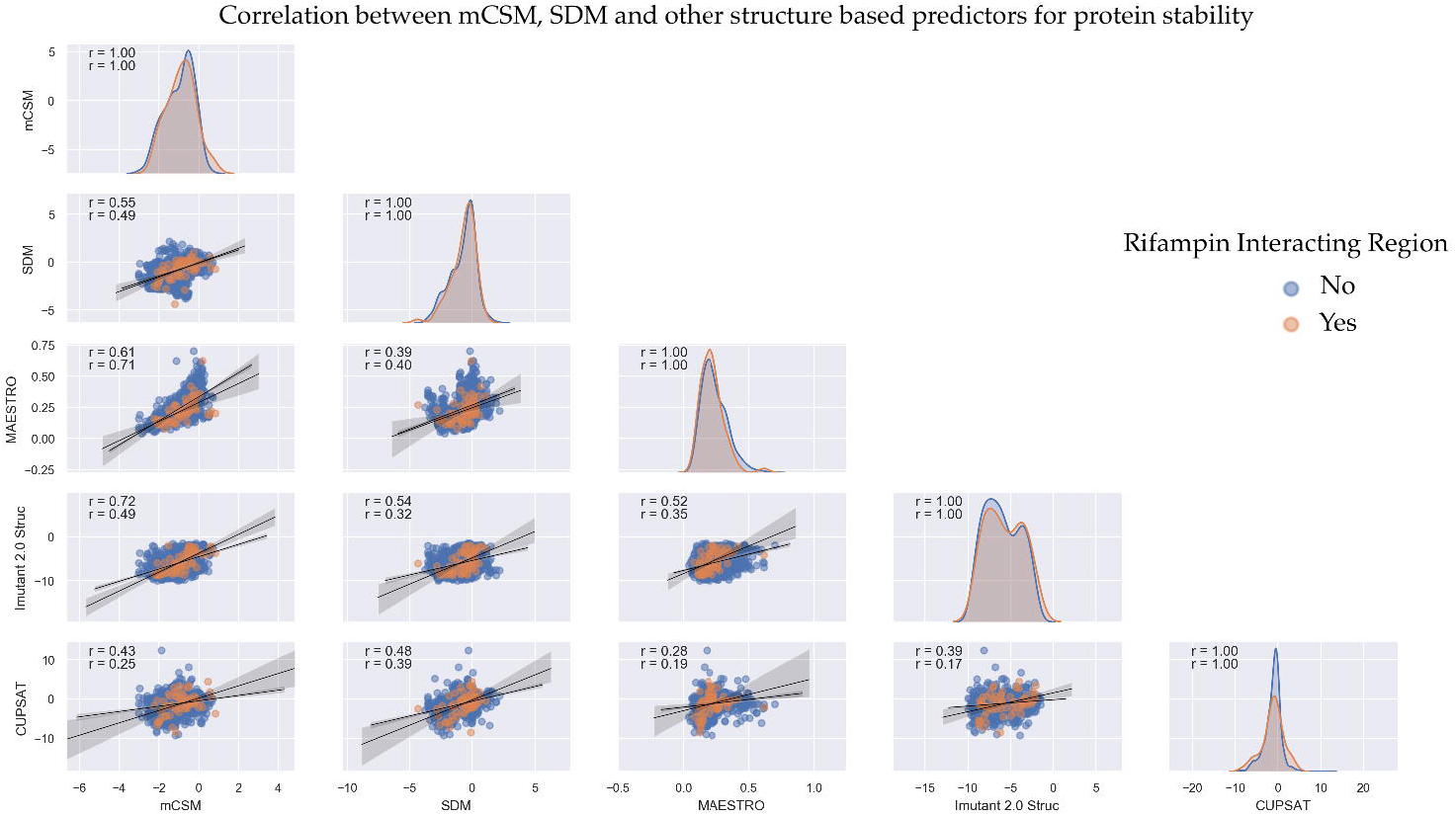
Pairplot depicting correlations between mCSM, SDM and other structural predictors of protein stability changes upon mutations in the β subunit of RNAP. Each datapoint corresponds to maximum destabilizing effect noted at each residue position in the β subunit when systematically mutated to other 19 residues. The data points in orange correspond to predictions at rifampin interacting residues.

**Supplementary Figure 2:**
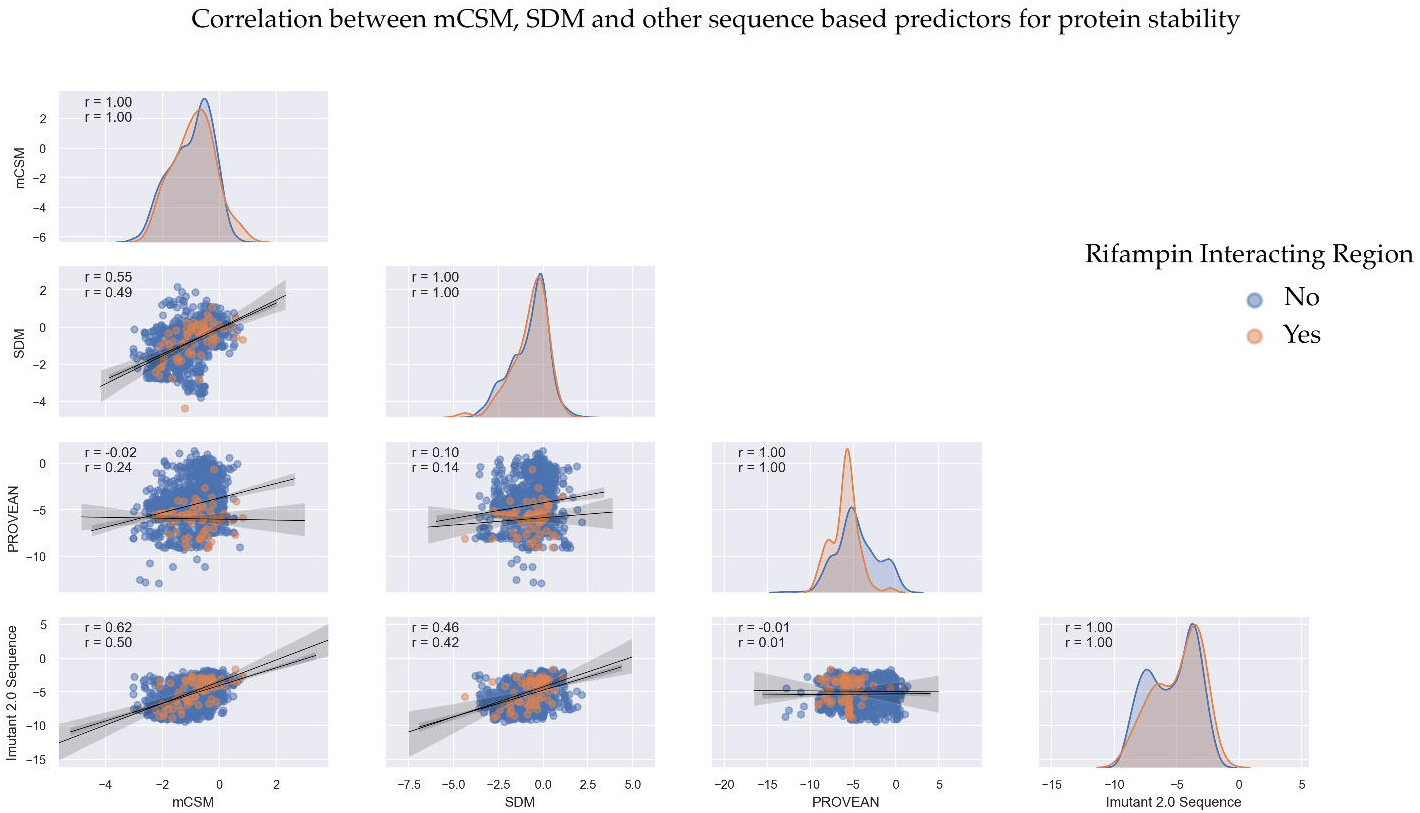
Pairplot depicting correlations between mCSM, SDM and other sequence-based predictors of protein stability changes upon mutations in the β subunit of RNAP. Each data point corresponds to maximum destabilizing effect noted at each residue position in the β subunit when systematically mutated to other 19 residues. The data points in orange correspond to predictions at rifampin interacting residues.

**Supplementary Figure 3:**
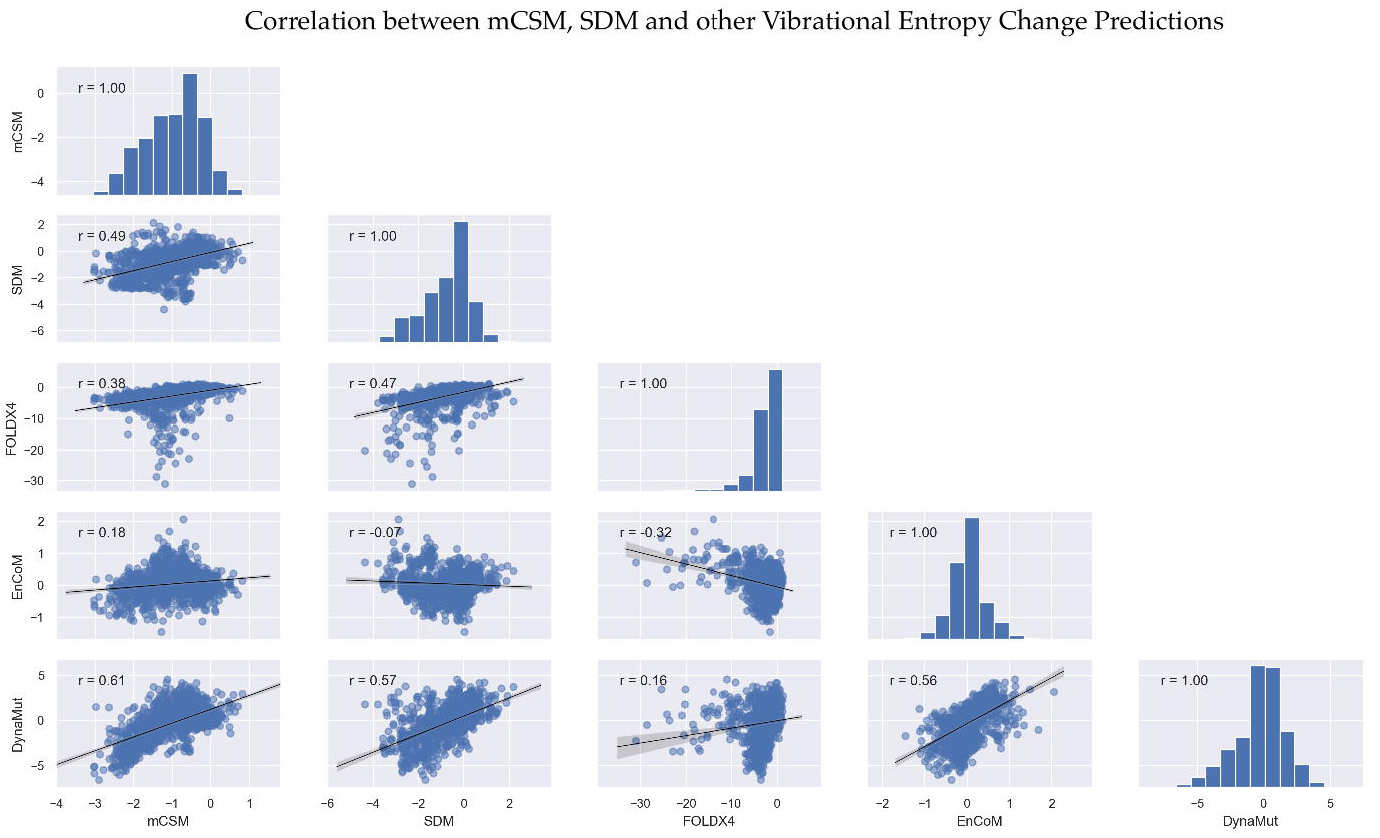
Pairplot depicting correlations between mCSM, SDM and other NMA-based predictors of protein stability changes upon mutations in the β subunit of RNAP. Each data point corresponds to average destabilizing effect noted at each residue position in the β subunit when systematically mutated to other 19 residues.

